# PIEZO1 and PIEZO2 Blade domains are differentially required for channel localization and function

**DOI:** 10.1101/2024.04.06.588398

**Authors:** Sergio Sarrió-Ferrández, Espe Selva, Francisco J Taberner

## Abstract

PIEZO1 and PIEZO2 are critical force-gated ion channels, detecting and transducing mechanical forces into ionic currents in many eukaryotic cell types, serving essential physiological roles. CryoEM and structure-function studies have revealed that three PIEZO monomers assemble as a 3-blade propeller, highlighting essential structural aspects for channel function. One of the most prominent features in PIEZO architecture is the Blade, a large membrane embedded domain that comprises 36 transmembrane fragments organized in 9 THU (Transmembrane Helix Units). Despite its suggested role in force transduction, the contribution of the Blade domain in channel physiology remains unclear. By systematically generating different truncated versions of PIEZO1 and PIEZO2, lacking parts of the Blade, we show the intact PIEZO1 Blade is essential for proper localization and function. Conversely, our results indicate the PIEZO2 Distal Blade segments (THU1-3) are dispensable for normal mechanical sensitivity. However, it plays a central role in channel stability and localization, containing a region that mediates the intracellular retention of a chimeric membrane protein. Our study indicates that, in addition to their biophysical properties, PIEZO1 and PIEZO2 also differ in the regulation of their localization, adding a new layer of control on PIEZO2 activity.

## Introduction

Sensing and responding to environmental cues is crucial for the survival of organisms. Among the diverse stimuli that animals encounter, mechanical forces are among the most prominent. In multicellular organisms, from worms to mammals, the ability to detect surrounding mechanical forces primarily relies on the two members of the PIEZO family of ion channels, PIEZO1 and PIEZO2 that respond to cell deformation evoking inactivating currents^1^.

Soon after its discovery, PIEZO1 was regarded as the major force transducer in non-neuronal cells where it also involved in sensing traction forces in motile or migrating cells, as well as in volume regulation in red blood cells^2–4^. PIEZO2 though, was recognized as the main transducer of mechanical cues in primary sensory neurons, playing a crucial role in touch and proprioception in mice and humans^5–10^. Notably, an ever-expanding number of studies are uncovering new functions for PIEZO channels, further highlighting their physiological relevance. For instance, recent studies show that PIEZO1 is critical for lymphatic valve development^11^. In the case of PIEZO2, recent work highlighted their important role in interoception, have pinpointed critical roles for PIEZO2 in sexual function, urination and respiratory control^11–13^ and underscored a significant role in pain transduction in humans and in rodents^14–16^

PIEZO channels are among the largest transmembrane proteins expressed in eukaryotic cells. They assemble as trimers, with each subunit exceeding 2,800 amino acids in length in the case of mouse PIEZO2. This substantial size, and the channel conformation, induce deformations in the cell membrane, visualized by cryo-electron microscopy (cryo-EM) as membrane invaginations^17, 18^. The cryo-EM structures of PIEZO1 and PIEZO2 reveal a common propeller-like architecture, with three blades emanating from a central cation-selective pore^18–21^. Each Blade domain is allosterically linked to the central pore through interdomain interactions involving the Beam (an extensive alpha helix located beneath the central third of the blade), the C-terminal domain (CTD), and the Latch domain. These interactions are crucial for fine-tuning the mechanical sensitivity of PIEZO to specific stimuli as well as limiting its unitary conductance^19, 22^.

A prominent feature of PIEZO channels is the extensive Blade domain, formed by nine consecutive groups of four transmembrane helices known as Transmembrane Helix Units (THUs). Each THU is connected to its neighbors by intracellular loops lacking a defined structural conformation, also known as intrinsically disordered regions (IDRs). It has been recently shown that the IDRs contribute to some of the biophysical differences observed between PIEZO1 and PIEZO2 currents, such as velocity dependence of current amplitude, limited stretch response, and cytoskeleton-mediated activation in PIEZO2^23^. The function of the Blade is however still obscure. Despite evidence suggesting a central role for the Blades in sensing membrane deformation^19, 24^, the impact of the Blade’s THUs on mechanical responses and other aspects of PIEZO channel cellular physiology, such as protein stability and subcellular localization, remain largely unexplored.

Here, we have investigated the role of PIEZO Blade domain by generating a series of PIEZO1 and PIEZO2 N-terminal truncated variants lacking distal THUs, and analyzed their roles using electrophysiology, super-resolution microscopy and analysis of protein stability assays. We found that PIEZO2 Distal Blade region encompassing THU1-3 are totally dispensable for channel conducting activity with minimal variations in their biophysical properties. However, this region regulates channel localization and stability. Importantly, the same region in PIEZO1 is required for proper membrane localization and function. The marked difference in subcellular localization of equivalent PIEZO1 and PIEZO2 mutants suggests that PIEZO2 possesses an additional mechanism controlling the channel membrane expression that is not present in PIEZO1.

## Results

### Distal Blade is required for PIEZO1 pocking-evoked currents but is dispensable for PIEZO2 activity

PIEZO2 Cryo-EM structure reveals nine well-defined Transmembrane Helix Unit domains (THUs; THU1-THU9) conforming the Blade domain^21^. In contrast, the different PIEZO1 Cryo-EM models have only solved THUs 4-9^18–20^, likely indicating that the missing THU1-3 are highly mobile within the membrane, consequently hindering their structural resolution^25^. This structural divergence between PIEZO1 and PIEZO2 Blade domain suggest that it could have different impact in channel physiology. To investigate the Blade domain’s roles, and to address potential differences between PIEZO1 and PIEZO2, we conceptually divided the Blade into three regions (Fig 1A). The Distal Blade (THU1-3) encompasses the THUs furthest from the channel pore which in PIEZO1 are more plastic that in PIEZO2. The Medial Blade (THU4-6) comprises three THUs with extracellular loops implicated in mechanosensitivity^19^. Finally, the Central Blade (THU7-THU9) that is closely associated with critical gating structures such as the CTD, Beam, and Anchor^19, 20^.

**Fig. 1.**
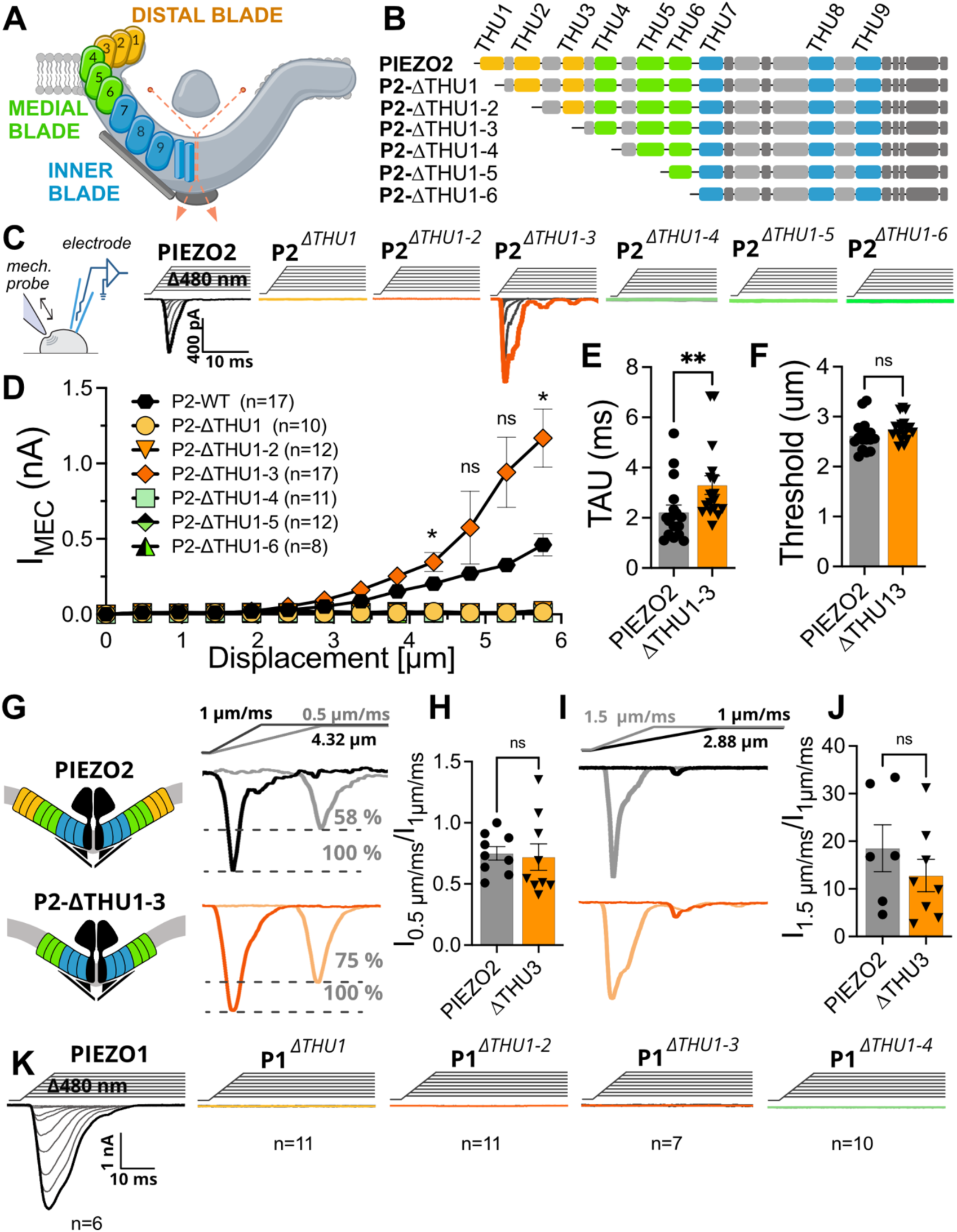
Truncations of PIEZOs Blade Domain alters indentation evoked currents. **A)** Diagram of the regions of the PIEZO2 Blade Domain. **B)** Schematic representation of different PIEZO2 truncated mutants. **C)** Sketch of whole-cell mechano-clamp recording and whole-cell example traces of PIEZO2 WT and the truncated versions expressed in N2a-P1KO cells. **D)** Displacement-responses curves of peak current amplitudes of PIEZO2 and the deletion mutants. Symbols represent mean±s.e.m., (n≥8, two-way ANOVA). **E)** Comparison of the inactivation time constant of PIEZO2 WT and PIEZO2-ΔTHU1-3 truncated version, (n= 17, t-test). **F)** Comparison of the mechanical threshold of PIEZO2 WT and PIEZO2-ΔTHU1-3 truncated versions, (n= 17, t-test). **G)** Example traces of PIEZO2 WT and PIEZO2-ΔTHU1-3 truncated version at 1 µm/ms (black and orange) and 0.5 µm/ms (gray and pale orange) indentation velocity. **H)** Comparison of current amplitude ratios at slower indentation velocities (0.5µm/ms/1µm/ms) of wild-type and P2-ΔTHU1-3 channels (n=9, t-test). **I)** Example traces of PIEZO2 WT and PIEZO2-ΔTHU1-3 truncated version at 1 µm/ms (black and orange) and 1.5 µm/ms (gray and pale orange). **J)** Comparison of current amplitude ratios at higher indentation velocities (1.5µm/ms/1µm/ms) of wild-type and P2-ΔTHU1-3 channels (n≥ 6, t-test). **K)** Example traces of whole-cell PIEZO1 and truncated versions currents.

To address the functions of the Blade domain, we sequentially trimmed the Distal and Medial THUs, creating a set of truncated PIEZO2 versions (Fig 1B). We then evaluated their function in Neuro2a-PIEZO1-KO cells, which lack endogenous PIEZO1 and exhibit no mechanically activated currents^24^. We initially compared currents evoked with mechanical indentation of the cell membrane (pocking) by whole-cell patch-clamp (Fig 1C). As shown in Figures 1C and 1D, deletion of THU1 alone (PIEZO2-ΔTHU) or from THU1 to THU2 (PIEZO2-ΔTHU1-2) rendered the channel unresponsive to mechanical stimuli. Unexpectedly, cells expressing the truncated PIEZO2 lacking the entire Distal Blade (PIEZO2-ΔTHU1-3) did respond to membrane indentation. Indeed, these cells displayed significantly increased mechanically activated currents at high indentations (PIEZO2: 0.46 ± 0.07 nA vs PIEZO2-ΔTHU1-3: 1.16 ± 0.38 nA at 5.8 µm indentation, Fig 1C, D). PIEZO2-ΔTHU1-3 currents showed a similar activation threshold to the wildtype channel (threshold: PIEZO2: 2.26 ± 0.30 µm vs PIEZO2-Δ1-3: 2.75 ± 0.23 µm, Fig 1F) but a slight, yet significant, increase in the inactivation time constant (τ: PIEZO2: 2.23 ± 0.29 ms vs. PIEZO2-Δ1-3: 3.30 ± 0.37 ms, Fig 1E). Noteworthy, this activity was completely lost with further THU deletions of the Medial Blade indicating that. while the Medial Blade is required for channel activity, PIEZO2 Distal Blade is dispensable.

To further characterize the effect of the Distal Blade deletion on the properties of PIEZO2 macroscopic currents, we studied the PIEZO2 well-documented velocity dependence of the peak current amplitude. We therefore compared the maximal current intensities at slower (0.5 μm/s) and faster (1.5 μm/s) indentation velocities. We observed that, reducing the indentation velocity from 1 μm/s to 0.5 μm/s in the PIEZO2-ΔTHU1-3 mutant resulted in a similar decrease in maximal current amplitude (0.75 ± 0.05 fold reduction) as in the wildtype channel (0.71 ± 0.11 fold reduction, Fig 1G). Analogously, increasing the indentation velocity from 1 μm/s to 1.5 μm/s resulted in larger mechanical currents at smaller indentations (2.88 µm) with no significant difference in the fold change between PIEZO2 and P2-ΔTHU1-3 (fold change: PIEZO2: 18.54 ± 4.93 vs P2-ΔTHU1-3: 12.80 ± 3.42, Fig 1I). These results indicate that the Distal Blade is not involved in the PIEZO2 enhanced current velocity dependence, consistent with the role of IDR2 in regulating this property^23^. Hence, the subtle differences in the overall biophysical properties between wildtype PIEZO2 and the truncated version lacking the Distal Blade domain (except for the increased current amplitude) indicate that this part of the channel is not required for detecting indentation forces nor to regulate mechanical sensitivity.

We next evaluated the role of the Distal Blade in PIEZO1 by undertaking a similar approach of domain trimming. To this end, we sequentially deleted the protein sequence from the beginning to the end THU1, from THU1 to THU2, from THU1 to THU3 and from THU1 to THU4, which include the Distal Blade and the first THU of the Medial Blade. In marked contrast to the results obtained in wildtype PIEZO1 where all the patched cells responded with currents to cell pocking (6 out of 6), we did not detect any mechanically activated currents in response to the same stimulus in cells expressing any of truncated PIEZO1 versions (0 out of 10, for each of the indicated mutants, Fig 1K). Therefore, these results demonstrate that, in contrast to PIEZO2, PIEZO1 Distal Blade is essential for the cells to respond to membrane deformation by indentation.

### PIEZO2 Distal Blade is not required for channel stretch sensitivity

Considering that PIEZO2 mechanical activation involves two relatively independent force transmission mechanisms (force from membrane and force from filament^23^), the ability of the Distal Blade truncation (PIEZO2-ΔTHU1-3) to respond to cell pocking suggests the filament mechanism remains intact. However, this doesn’t necessarily preclude the force from membrane mechanism being affected. Therefore, we investigated if shortening the PIEZO2 Distal Blade domain, consequently reducing the Blade-membrane interaction surface, affects stretch-induced responses. To this end, we recorded stretch-activated currents in cell-attached configuration by applying incremental negative pressure steps to the membrane patch in cells expressing PIEZO2 and the three Distal Blade truncations. As reported previously^23^, only a fraction (34.8 %) of PIEZO2-expressing cells displayed stretch-evoked currents (Fig 2A, B). Conversely, no stretch currents were recorded in any cells expressing PIEZO2-ΔTHU1 (0 out of 10 cells) or PIEZO2-ΔTHU1-2 (0 out of 10 cells, Fig 2B). Strikingly thought, removing the complete Distal Blade (PIEZO2-ΔTHU1-3) rendered cells with stretch currents in a similar proportion to the wildtype channel (36.8 %, Fig 2a and 2b).

**Fig 2.**
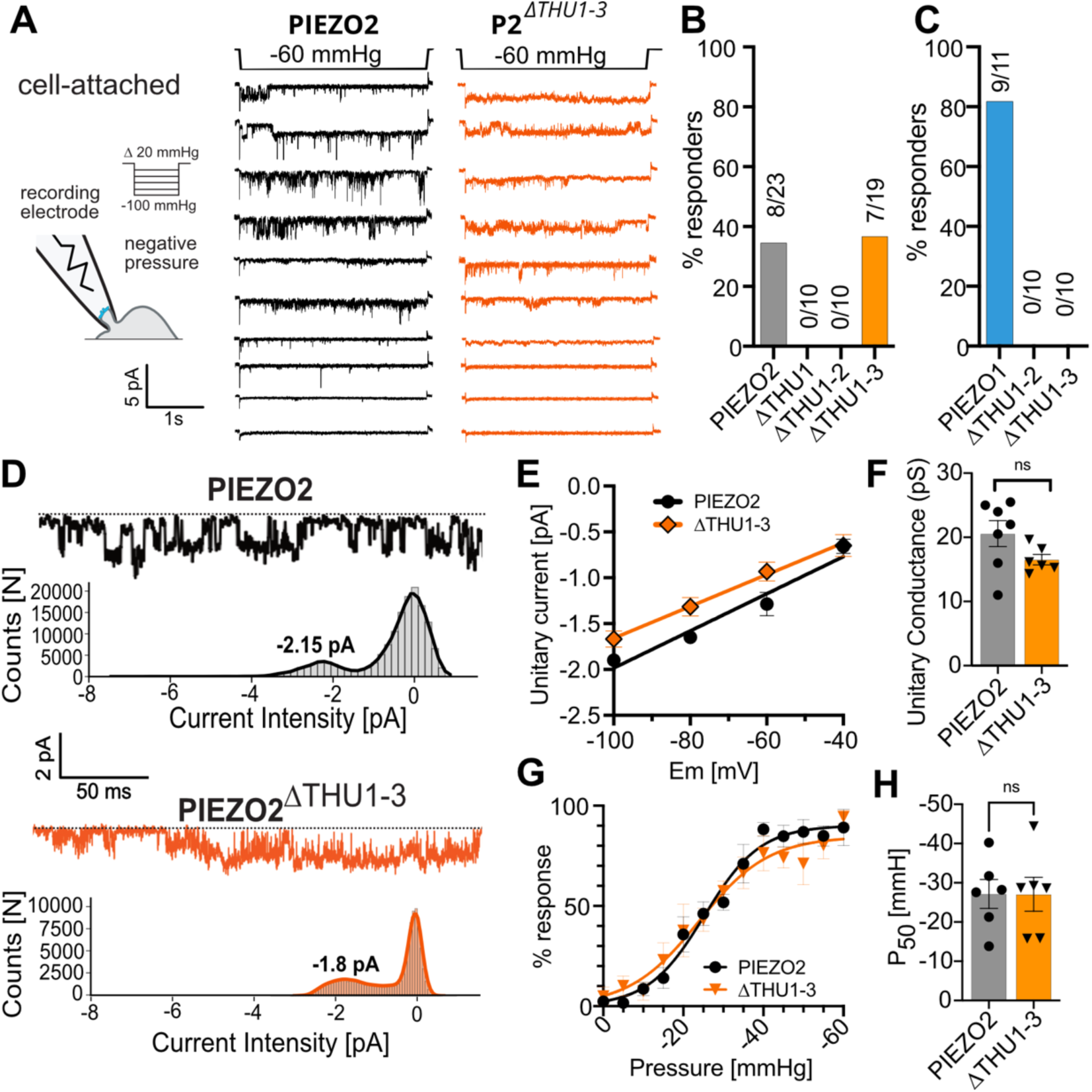
The Distal Blade deletion mutant does not affect PIEZO2 stretch activated responses. **A)** Sketch of Single-Channel (Pressure-Clamp) recordings configuration, and example traces of stretch-activated PIEZO2 WT and PIEZO2-ΔTHU1-3 currents recorded at −60 mV and a pressure stimulus of −60 mmHg. **B)** Comparison of percentage of stretch responders PIEZO2 and the Blade truncated versions (n≥10). **C)** Comparison of percentage of stretch responders PIEZO1 and the Blade truncated versions (n≥10). **D)** Example traces (top) of stretch-activated PIEZO2 (black), PIEZO2-ΔTHU1-3 (orange) are shown together with corresponding current amplitude distribution histograms. **E)** Linear regression fits of the I-V curves of the responder channels. Symbols represent mean±s.e.m. (n≥6). **F)** Comparison of the Unitary Conductance of the responders’ channels, WT (gray) and ΔTHU1-3 (orange). The bar graphs represent mean±s.e.m. (n≥6, t-test). **G)** Pressure-Response curve of PIEZO2 and PIEZO2-ΔTHU1-3 (n=6). Symbols represent the AUC normalized to the maximum value. **H)** P50 of PIEZO2 WT and PIEZO2-ΔTHU1-3 truncated version. The bar graph represents as mean±s.e.m. the pressure in mmHg at which the response is half maximum (n=6, t-test).

We next compared stretch responses between PIEZO1, PIEZO1-ΔTHU1-2 and PIEZO1 ΔTHU1-3 deletion mutants. As shown in Fig 2C, while a high proportion of PIEZO1-expressing cells responded to membrane stretch (9 out of 11), no activity was observed in cells expressing the PIEZO1 truncates (0 out of 10 in both assayed mutants). Thus, while the Distal Blade is required for PIEZO1 stretch responsiveness, it is unnecessary for PIEZO2 to detect membrane tension, suggesting that the extent of membrane-channel interaction surface is not critical in PIEZO2.

The fact that there is virtually no difference in the percentage of stretch responders of the PIEZO2-ΔTHU1-3 truncate when compared to wildtype PIEZO2, does not exclude the possibility that the current properties of the stretch-evoked responses are different. Therefore, we further characterized some biophysical properties of the PIEZO2-ΔTHU1-3 mutant currents. When analyzing stretch-activated single-channel openings of the Distal Blade truncated PIEZO2, we observed that they were slightly smaller than the wildtype ones at all recorded voltages (unitary current at −100 mV; PIEZO2: 1.90 ± 0.21 pA vs PIEZO2-ΔTHU1-3: 1.68 ± 0.21 pA; Fig 2D, E). Consequently, this resulted in a reduced, but not statistically significant unitary conductance (PIEZO2: 20.57± 2.01 pS vs PIEZO2-ΔTHU1-3: 16.53 ± 0.82 pS; Fig 2F). Hence, as expected, a mutation located far away from the pore has marginal effects on the channel conductance.

To assess whether the deletion of the Distal Blade alters channel’s sensitivity to membrane stretch, we next compared the pressure sensitivity of the mutant and wildtype channel by measuring the charge transfer (Area Under the Curve) at different pressures. As shown in Fig 2G, both channels displayed near overlapping pressure sensitivity curves, resulting in practically identical half-activation pressures (P_50_, PIEZO2: −27.15 ± 3.68 mm Hg vs PIEZO2-ΔTHU1-3: −27.08 ± 4.33 mm Hg; Fig 2H). These biophysical results indicate that stretch-activated responses are virtually identical between PIEZO2 wildtype and the PIEZO2-ΔTHU1-3 deletion mutant. Therefore, these findings evidence that in PIEZO2, 1/3^rd^ of the Blade domain is irrelevant for channel’s electrical activity. Noteworthy, the different effect of the Distal Blade removal in PIEZO1 and PIEZO2 strongly suggests a distinctive role of this region in the physiology of each PIEZO channel.

### The Distal Blade regulates PIEZO2 stability

Although PIEZO2 Distal Blade is dispensable for channel function and mechanical sensitivity, the fact that partial Distal Blade truncations in PIEZO2 rendered a channel unable to respond to membrane indentation suggests that these initial THUs are relevant for PIEZO2’s physiology. Given the lack of pocking and stretch responses in PIEZO2-ΔTHU1 and PIEZO2-ΔTHU1-2, we initially hypothesized that these trimmed PIEZO2 versions might have defects in protein expression. To test this hypothesis, we examined protein levels of the mutants in total extracts from transiently transfected Neuro2a-PIEZO1-KO using Western Blotting (Fig 3A). When normalized to the β3-tubulin loading control (Fig 3B), PIEZO2-ΔTHU1 showed a consistent, but not statistically significant, decrease in protein levels (fold change compared to wild type; PIEZO2-ΔTHU1: 0.49 ± 0.10). In contrast, PIEZO2-ΔTHU1-2 expression was enhanced but did not differ from wild type significantly (PIEZO2-ΔTHU1-2: 2.15 ± 0.73). Noteworthy, the Distal Blade deletion mutant (PIEZO2-ΔTHU1-3) displayed clearly higher protein levels than the wildtype (PIEZO2-ΔTHU1-3: 5.89 ± 1.71). Therefore, alterations in protein levels do not explain the lack of mechanical responses in the PIEZO2-ΔTHU1 and ΔTHU1-2 truncated mutants, but likely account for the increased macroscopic current observed in the PIEZO2-ΔTHU1-3 mutant compared to wildtype (Fig 1C and 1D).

**Fig 3.**
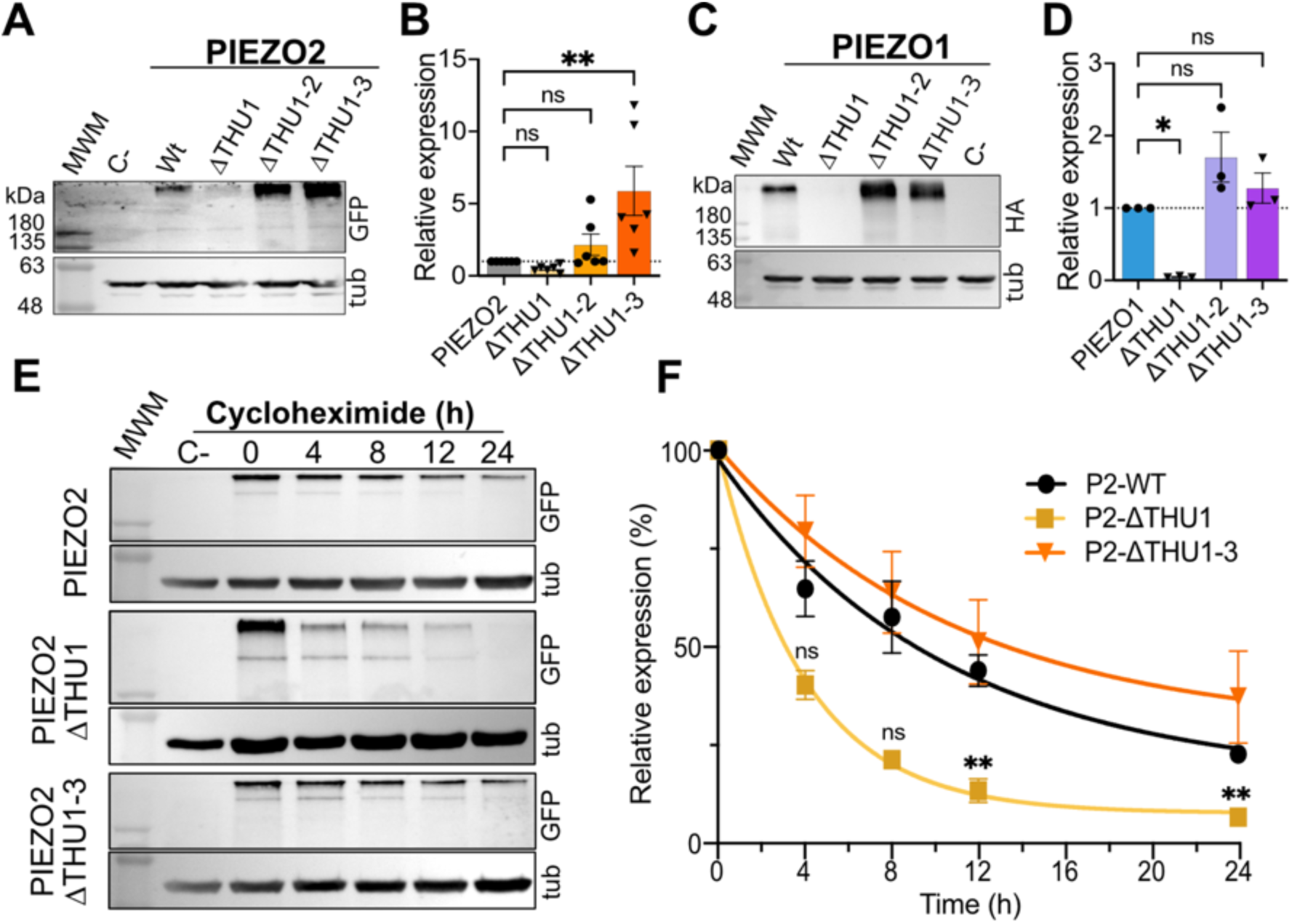
PIEZO1 and PIEZO2 Distal Blade deletion modify channel expression and stability. **A)** Western blot of whole-cell lysate of PIEZO2 and the PIEZO2 truncated versions expressed in N2a-P1-KO cells. **B)** Relative protein expression of PIEZO2 truncated variants, the bar graph shows the relative expression levels normalized to the WT signal as mean±s.e.m., (n=6, One-way ANOVA). **C)** Western blot of whole-cell lysate of PIEZO1 and the PIEZO1 truncated versions expressed in N2a-P1-KO cells. **D)** Relative protein expression of PIEZO1 truncated variants, the bar graph shows the relative expression levels normalized to the WT signal as mean±s.e.m., (n=3, One-way ANOVA). **E)** Cycloheximide protein stability assay of PIEZO2 WT, ΔTHU1 and ΔTHU1-3 truncated versions, **D)** Stability plots of PIEZO2 WT, ΔTHU1 and ΔTHU1-3 truncated versions, symbols represent the relative expression levels normalized to the T0 for each channel at all timepoints measured as mean±s.e.m., (n≥3, Two-way ANOVA).

Using the same methodology, we next examined the expression levels of truncated PIEZO1 variants lacking the Distal Blade domain. Consistent with our PIEZO2 observations, quantification and normalization of the signal revealed a significant reduction in PIEZO1-ΔTHU1 protein levels (fold change compared to wild type; PIEZO1-ΔTHU1: 0.05 ± 0.01, Fig 3C). Conversely, total expression of PIEZO1-ΔTHU1-2 and PIEZO1-ΔTHU1-3 showed a modest non-significant increase. (PIEZO1-ΔTHU1-2: 1.70 ± 0.35; PIEZO1-ΔTHU1-3: 1.28 ± 0.21, Fig 3C). Therefore, as for PIEZO2, variations in protein levels do not explain the absence of mechanical responses. Interestingly though, the reduction in protein amount in both PIEZOs upon THU1 deletion suggests a shared role for this region in regulating channel expression levels.

Cellular protein abundance is determined by the equilibrium between protein synthesis and degradation. Considering that all PIEZO variants are expressed from the same promoter and that protein degradation is more frequently regulated than protein synthesis, we hypothesized that the rate of protein degradation might be altered in the PIEZO2-ΔTHU1 and PIEZO2-ΔTHU1-3 mutants. To test this hypothesis, we inhibited protein synthesis with cycloheximide, a translation elongation blocker in cells expressing the wildtype and the mutant channels. We then monitored protein levels over a 24-hour period (Fig 3E) after cycloheximide addition. The abundance of wildtype PIEZO2 decreased to 22.33 ± 2.03% within 24 hours, with a half-life of 7.21h hours estimated from the monoexponentially decay fitted data. The more abundant PIEZO2-ΔTHU1-3 mutant displayed a slight increase in stability, retaining 37.0 ± 1.2% of its expression at 24 hours with an estimated half-life of 7.57 hours (Fig 3F). Interestingly, the degradation of PIEZO2-ΔTHU1 was faster, with only 6.3 ± 2.3% of the protein remaining after 24 hours and an estimated half-life of 2.77 hours. These findings suggest that the Distal Blade region of the PIEZO channels directly regulates channel stability. Additionally, they point towards a potential regulatory mechanism involving the THU2-THU3 region. This region appears to harbor a degradation signal that can be masked only if THU1 is present, suggesting the possibility that proteins interacting through THU1 may regulate PIEZO channels abundance.

### PIEZO2 Distal Blade regulates subcellular channel distribution

Given that the protein levels of non-functional PIEZO2-ΔTHU1 and PIEZO2-ΔTHU1-2 mutants are unaltered, we investigated if the hindered function was due to deficient protein trafficking. To test this tenet, we co-transfected Neuro2a-PIEZO1-KO cells with the different PIEZO2 mutants and the Endoplasmic Reticulum marker protein ER-mCherry (mCherry red fluorescent protein fused to the Chicken Lisozyme export sequence) and visualized both proteins by immunofluorescence followed by LSM880-Airyscan super-resolution imaging. As previously described^23, 26^, full PIEZO2 channels show a punctuated pattern distribution with minimal overlap with the ER (Fig 4A) as indicated by the low Pearson’s coefficient obtained (PIEZO2: 0.42 ± 0.04, Fig 4B). Notably, this punctate localization became reticular with a high degree of colocalization with the ER marker in the PIEZO2-ΔTHU1 and PIEZO2-ΔTHU1-2 mutants, with high Pearson’s coefficients of 0.81 ± 0.04 and 0.73 ± 0.07 respectively. This observation suggests that THU1 is required for proper PIEZO2 distribution, and its absence severely compromises channel exit from the ER. Interestingly, the complete deletion of the Distal Blade (PIEZO2-ΔTHU1-3) restored the punctate wildtype pattern with similar punctum size (Punctum diameter, PIEZO2 0.59 ± 0.05 µm, PIEZO2-ΔTHU1-3 0.70 ± 0.05 µm, Fig 4C). This pattern change was accompanied by a decrease in ER colocalization indicated by a reduction in the Pearson’s colocalization coefficient. However, the Pearson’s coefficient (0.54 ± 0.03) did not reach wildtype values since there was still some faint reticular PIEZO2 signal likely due to the higher expression levels in this mutant. These results indicate that PIEZO2 Distal Blade is dispensable for proper channel localization.

**Fig 4.**
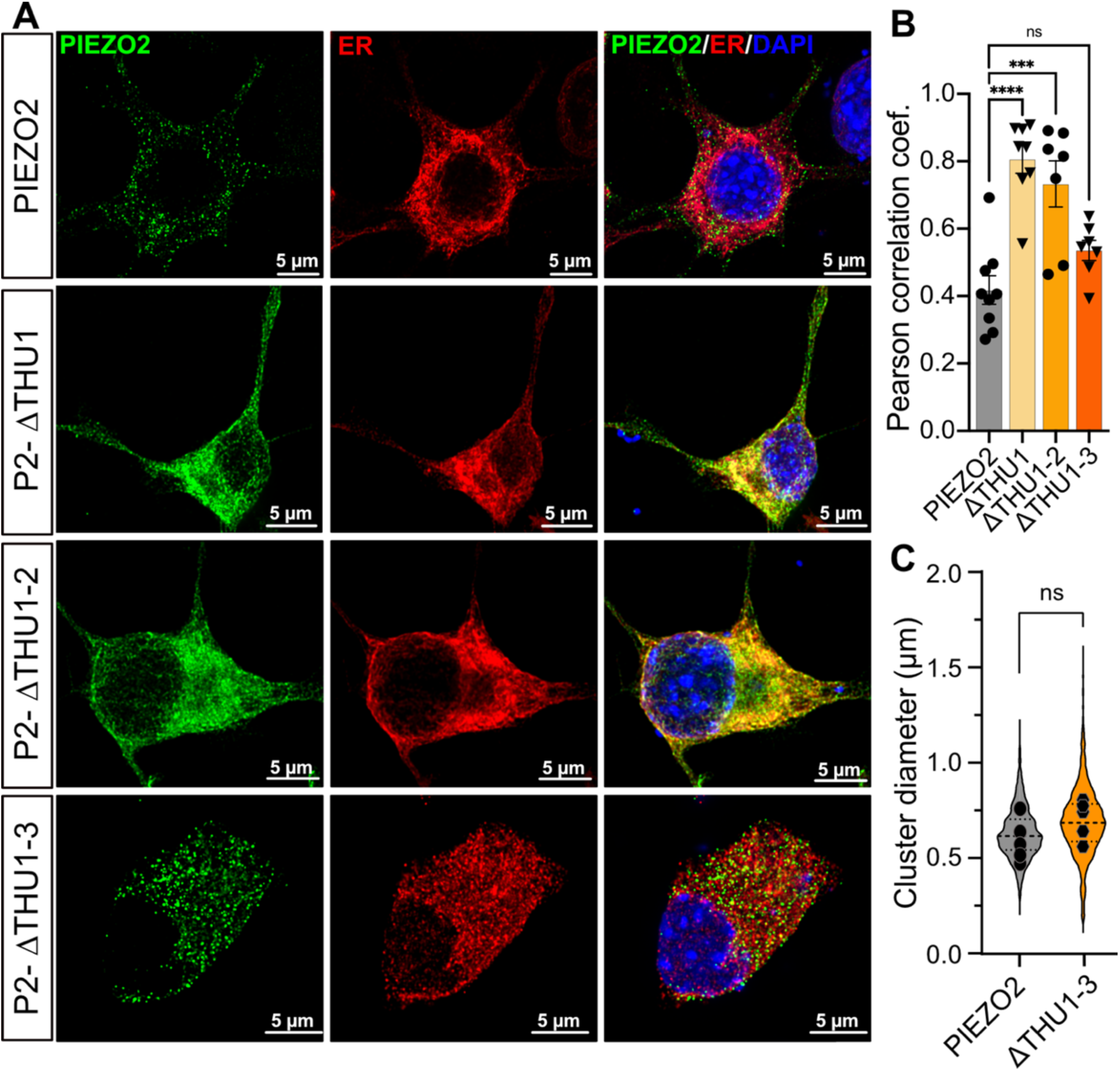
Distal Blade truncations differentially affect PIEZO2 subcellular localization. **A)** Immunocytochemistry of PIEZO2 WT and all truncated variants (anti-GFP antibody) along with an Endoplasmic Reticulum (ER) marker (anti-RFP antibody). **B)** Comparison of the Pearson correlation coefficient between each PIEZO2 variants and the ER. The bar graph shows the Pearson correlation coefficient as mean±s.e.m. (N= ≥7, One-way ANOVA). **C)** Comparison of the cluster diameter between P2 WT and the ΔTHU1-3 variant. Min-max violin plot showing the distribution of all cluster diameter treated independently (clusters WT:1340, P2-ΔTHU1-3: 1636). Each symbol corresponds with the mean diameter size per cell. The statistical analysis was done by comparing the mean of each cell (n=5, t-test).

We next studied the subcellular distribution of PIEZO1 Distal Blade mutants. As observed in Figure 5, PIEZO1 exhibited a punctuated pattern with an evident colocalization with the membrane marker Wheat Germ Agglutinin (WGA). Similarly to the observations in PIEZO2, PIEZO1 versions lacking the THU1 and THU1-2, localization pattern changed dramatically acquiring a reticulated pattern. Importantly, in marked contrast to PIEZO2, the deletion of PIEZO1 Distal Blade (ΔTHU1-3) did not recover the wild-type subcellular distribution, indicating that proper PIEZO1 localization requires the intact Blade Domain.

**Fig 5.**
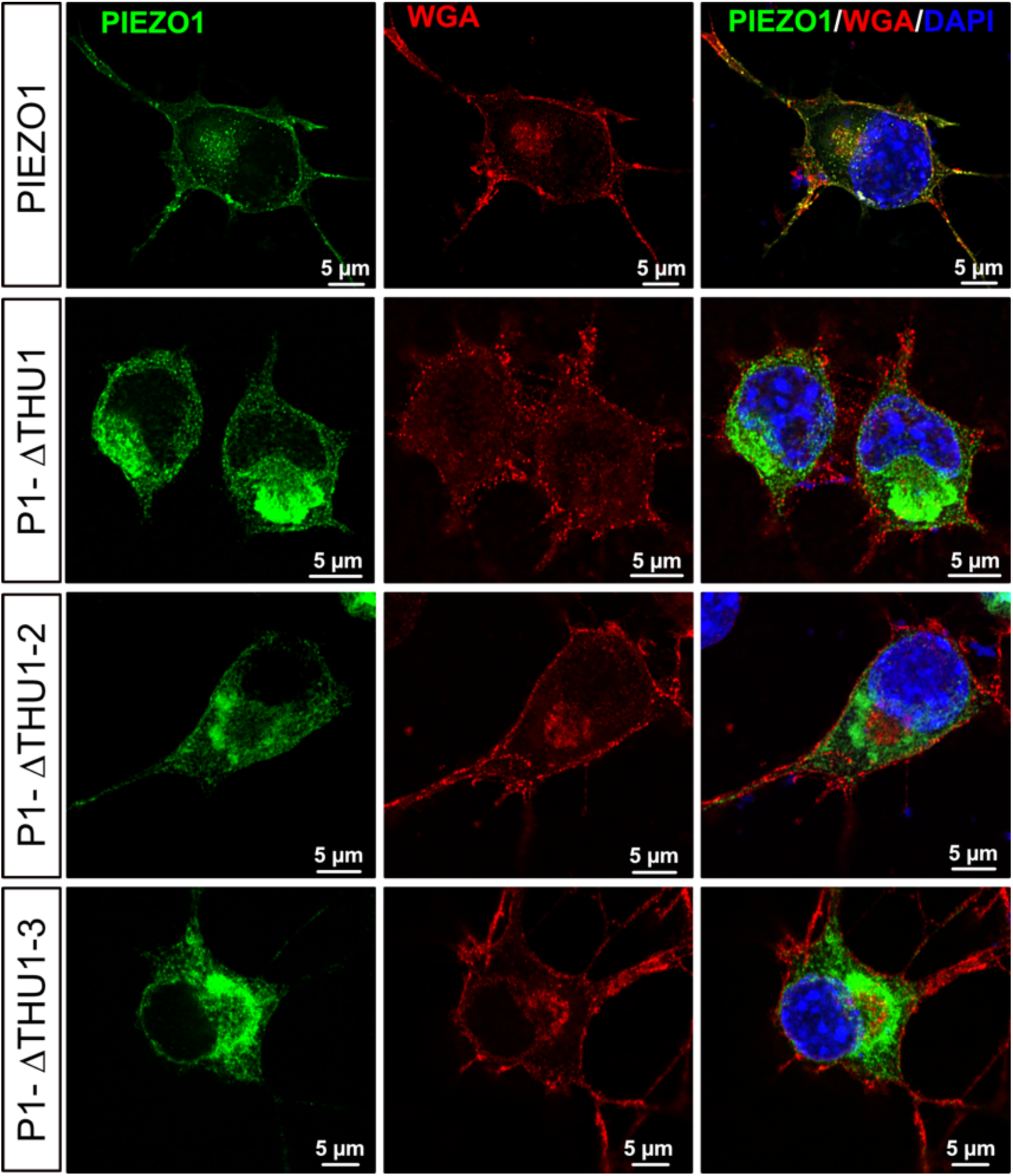
Subcellular localization of PIEZO1 and Distal Blade truncated variants. Immunocytochemistry of PIEZO1, PIEZO1-ΔTHU1, PIEZO1-ΔTHU1-2, PIEZO1-ΔTHU1-3 truncated variants (anti-GFP antibody) along with WGA Alexa 647 conjugated (Plasma membrane marker). Super-resolution images reveal that regular PIEZO1 punctuated subcellular distribution change to a reticulated patern upon the deletion of THU1. Unlike in PIEZO2 this anomalous distribution is nor reverted by removing the entire Distal Blade.

Altogether, these findings indicate that PIEZO1 and PIEZO2 Distal Blades serve different roles in the channel subcellular distribution. In the case of PIEZO2, it is not required for proper channel localization; however, it contains sequences that may fine tune PIEZO2 subcellular distribution, advocating a role in regulating PIEZO2 trafficking. All these aspects, not shared by PIEZO1, suggest that PIEZO2 membrane availability can be tuned under specific conditions or in certain cell types.

### PIEZO2’s THU3 mediates intracellular channel retention

The regulation of subcellular protein distribution often relies on specific protein regions that function as trafficking or retention signals^27^. Therefore, we investigated the presence of plasma membrane localization and endoplasmic reticulum (ER) retention sequences within the Distal Blade domain of PIEZO2. To achieve this, we cloned and expressed the PIEZO2 fragment encompassing the entire Distal Blade (amino acids 1-644) in Neuro2a-PIEZO1-KO cells. As shown in Fig 6A, PIEZO2’s Distal Blade does not show an evident plasma membrane localization pattern but highly co-localizes with the ER stain, indicating an inability to exit the ER. This finding further supports the notion that the Distal Blade is not required for reaching the membrane, but rather it contains an ER retention signal.

**Fig 6.**
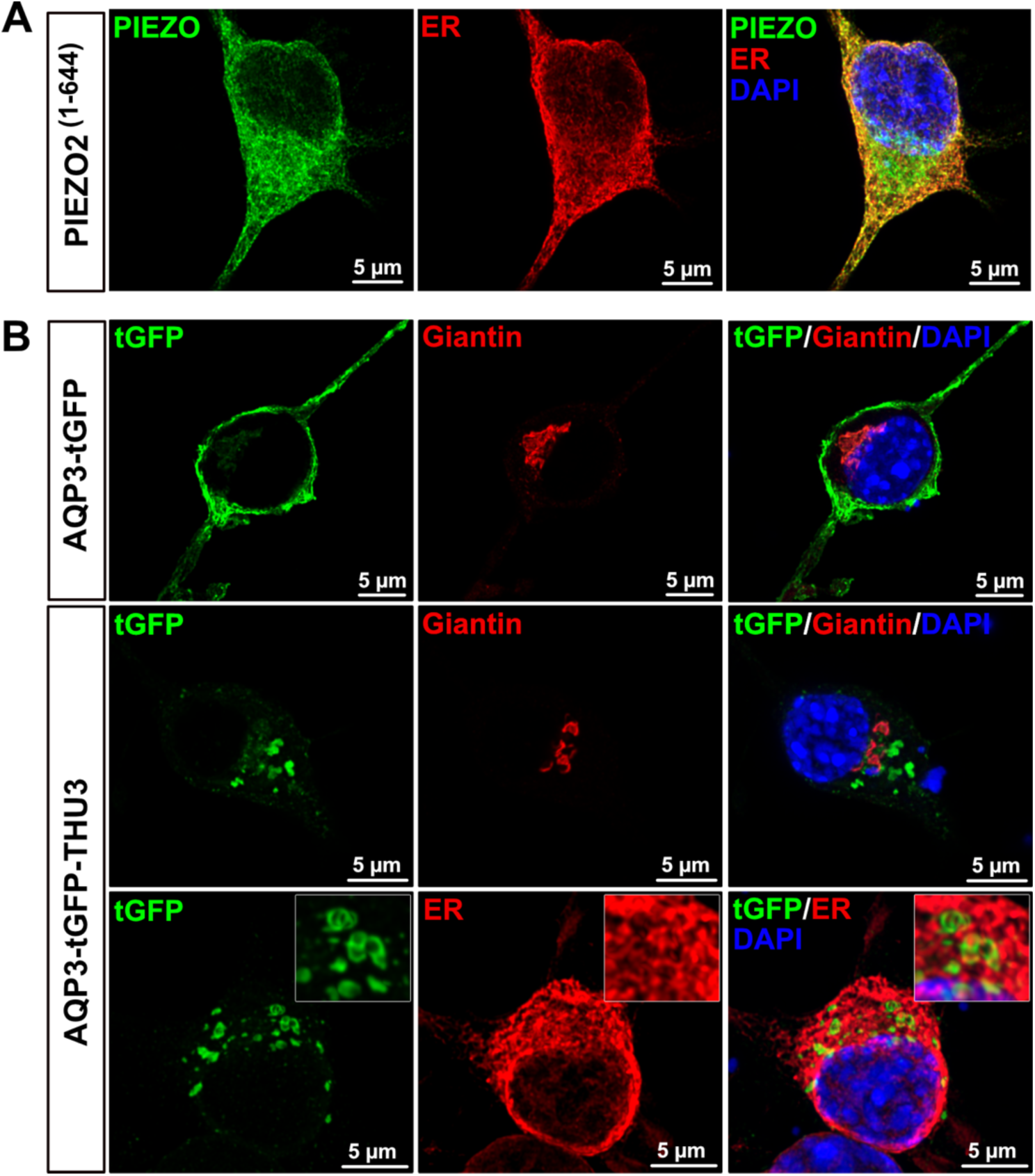
PIEZO2 THU3 mediates intracellular retention of a reporter membrane protein. **A)** Immunocytochemistry of PIEZO2**^(^**^1–644)^ encompassing the Distal Blade (anti-GFP antibody) along with an Endoplasmic Reticulum marker (ER-mCherry) (anti-RFP antibody). **B)** Immunocytochemistry of AQP3-tGFP (1st row) and AQP3-THU3-tGFP (anti-turboGFP antibody) along with cis Golgi Marker (2nd row, Giantin-mCherry) or an ER (ER-mCherry) (3rd row, anti-RFP antibody); the inset highlights the ring-shaped accumulations of the chimeric protein.

Since PIEZO2-ΔTHU1-2 is retained in the ER while PIEZO2-ΔTHU1-3 exhibits normal distribution, we hypothesized that the THU3 region might significantly contribute to ER retention. To test this hypothesis, we cloned PIEZO2 THU3 (amino acids 493-614) into the AQP3-tGFP membrane reporter system^28^. As expected, AQP3-tGFP displayed an evident membrane localization (Fig 6B). Interestingly, the AQP3-tGFP-P2-THU3 fusion protein exhibited a ring-shaped distribution (Fig 6B), often observed as granules in cells with higher expression levels. These structures did not colocalize with the cis-medial Golgi marker Giantin (Fig 6B). Notably, most AQP3-THU3 rings were located within the empty spaces of the ER network, with marginal peripheral overlap with the ER stain, resembling the characteristic ring-shaped structures of ER exit sites^17, 21, 22^. These results suggest that THU3 plays a role in PIEZO2 ER retention and further strengthen the idea that the Distal Blade is crucial for regulating the subcellular distribution of wild-type PIEZO2.

## Discussion

The overall aim of this study was to examine the role of the Blade domain in the physiology of PIEZO channels. We discovered three things: first, PIEZO1, but not PIEZO2 Blade, is required for proper channel function; second, Distal Blade (THU1-THU3) controls the PIEZOS stability; and third: PIEZO2 Distal Blade regulates the channel trafficking.

This domain is a salient feature in the PIEZO channel structure, not only by its large extension but also by the fact that most of it is embedded within the lipid bilayer^17, 18, 20, 21^. This fact makes the Blade ideally suited to detect membrane deformations caused by stimuli such as mechanical indentation or stretch. It has been described that some extracellular loops connecting transmembrane helices in PIEZO1 THU4 and THU5 are relevant in channel function as its deletion blocks stretch activated currents with some residual pocking activated currents^19^. Similarly, some of the intracellular loops connecting the THUs are also functionally relevant, conferring some of the distinctive functional properties between PIEZO1 and PIEZO2. A case in point being the role of the PIEZO2’s IDR2 in setting the characteristic velocity dependence of the channel which is relevant in mechanoreceptor function^23^. However, the relevance of the Blade length in channel responsiveness to stimuli has been not addresses. In addition to the function, it is plausible that this large domain also affects other aspects of PIEZO physiology as for instance the stability or the localization, properties that have been largely overlooked. So, with the aim of better understanding the role of this domain we incrementally trimmed PIEZO2 and PIEZO1 based on the THUs boundaries and compared the effect of the deletions on PIEZO1 and PIEZO2 mechanical sensitivity, stability, and localization.

Our study demonstrates that PIEZO1 and PIEZO2 Blade domains serve distinct functions for channel physiology. A key finding is that the PIEZO2 Distal Blade domain (composed of THU1-3 and corresponding intra- and extracellular loops) is not required for mechanically induced channel responses. Indeed, the PIEZO2 Distal Blade deletion mutant (PIEZO2-ΔTHU1-3) has minimal effects on the properties of pocking- and stretch-evoked currents. The fact that pocking mechanical thresholds and pressure sensitivity (P50) in stretch currents are similar to wild-type values indicates that this region is not involved in setting PIEZO2 mechanical sensitivity. In stark contrast, removal of the equivalent region in PIEZO1 renders the channel completely inactive. Recent studies on PIEZO1 conformational changes using nanoscopic fluorescence imaging suggest high flexibility in the PIEZO1 distal blade, potentially important for force transmission mechanisms^25^. Although our results with shortened PIEZO1 variants appear to support this notion, caution is warranted as the observed defects likely reflect a trafficking problem to the plasma membrane. Importantly, the force sensitivity of the PIEZO2 Distal Blade mutant, similar to wildtype, clearly demonstrates that the size of the PIEZO2 Blade does not linearly affect the channel’s ability to detect membrane thinning or reduce the response to cell pocking. Notably, further sequential deletions of the Medial Blade completely abolishing channel responses to both pocking and stretch, suggesting that these regions contain critical structures for the “force from lipids” and “force from filaments” mechanisms acting on PIEZO2^23^.

Despite the opposite functional consequences of complete Distal Blade (ΔTHU1-3) deletion in PIEZO1 and PIEZO2, our protein expression data suggest shared regulatory mechanisms controlling their levels. The restoration of reduced protein expression in PIEZO1-ΔTHU1 and PIEZO2-ΔTHU1 mutants by additional deletion of THU2 strongly indicates that this region is crucial for channel stability, not a consequence of protein dysfunction. Notably, PIEZO2-ΔTHU1 stability is severely compromised. Interestingly, some sodium and TRP channels regulate stability through activity-mediated endocytosis followed by degradation^31, 32^. However, this seems unlikely for the unstable PIEZO mutants. Immunocytochemistry reveals an anomalous intracellular localization affecting plasma membrane trafficking. Specifically, PIEZO2-ΔTHU1 and PIEZO2-ΔTHU1-2 mutants colocalize with the endoplasmic reticulum (ER), suggesting retention that disrupts the characteristic punctate PIEZO2 pattern. ER accumulation is associated with regulated transport. Similarly misfolded, potentially toxic or aberrant proteins as for instance mutants of the HERG potassium channel^33, 34^, are also often retained at the ER and subsequently enter the Endoplasmic Reticulum-Associated Degradation^35^ (ERAD) pathway. It is therefore plausible that Blade mutants misfold and undergo degradation. However, if misfolding solely governed protein levels and stability, reduced expression would be expected in both mis-trafficked ΔTHU1 and ΔTHU1-2 mutants of PIEZO1 and PIEZO2. Conversely, experimental evidence demonstrates normal ΔTHU1-2 levels, contradicting this hypothesis. These findings rather suggest a regulated mechanism where the channel region between the end of THU1 and THU2 (amino acids 139-355) contains a degradation signal normally masked by THU1. Hence, as in the case of TRPV1 channel^36^, PIEZO protein levels might be adjusted by regulated degradation through protein interactions. Given that PIEZO1 and PIEZO2 overactivity can lead to detrimental effects like Hereditary Xerocytosis^37, 38^, Distal Arthrogryposis^39^, or joint contractures^40^, it seems to be crucial to regulate PIEZO channel abundance and this proposed regulatory mechanism could be critical for this purpose.

An additional important finding of this study is the different contribution of the Distal Blade to PIEZO1 and PIEZO2 trafficking. It is well described that PIEZO1 and PIEZO2 exhibit a punctuated distribution within the cell, which likely represents the tendency of PIEZO channels to form clusters^23, 26, 41^. We show that an intact Blade with the 9 THUs is necessary for PIEZO1 acquiring their functional localization, since the deletion of the THU1 abrogates regular channel localization, phenotype that is conserved in other Distal Blade deletion mutants (PIEZO1-ΔTHU1-2 and PIEZO1-ΔTHU1-3). Importantly, we show that this is not the case for PIEZO2. Actually, the PIEZO2 Distal Blade (PIEZO2-ΔTHU1-3) mutant acquires the regular punctuated pattern. Additionally, the inability of the Distal Blade to localize the tGFP fluorescent protein into the plasma membrane or the prototypical PIEZO puncta further supports that this portion of the Blade is not involved in default protein trafficking. Interestingly though, the partial Blade deletions PIEZO2-ΔTHU1 and PIEZO2-ΔTHU1-2 accumulate in the ER and this retention is rescued by the additional deletion of the protein region including the IDR6 and THU3. Since IDR6 deletion does not alter channel activity^23^, the observed effect is likely due to the protein sequence defining the THU3. These findings suggest that the Distal Blade, although not necessary, influence protein localization. This suggestion is supported by the fact that PIEZO2 THU3 is able to change the localization of the membrane reporter Aquaporin3-tGFP into granular ring-shaped structures that reside withing the ER network and do not colocalize with the cis-Golgi, resembling the ER-Exit Sites. The fact that in wildtype PIEZO2, we do not observe these structures, indicate that normally this retention signal is somehow masked by the THU1-THU2 protein region, thus posing the tenet of a mechanism that regulate the membrane availability of PIEZO2 through ER retention. Such as mechanism, would help to tightly control the membrane abundance of PIEZO2 in a cell type dependent manner, as has been described for the TMC1 protein of the hair cell mechano-electrical transduction (MET) ion channels^42^. Further studies need to be performed probe and characterize this potential regulatory mechanism.

## Methods

### DNA constructs and generation of PIEZO mutants

For this study, we generated the we generated the GreenLanter C-terminal tagged PIEZO2 version (PIEZO2-GL) obtained by subcloning the mGreenLantern moiety from the LifeAct-mGreenLantern plasmid^43^ into a PIEZO2-HA-IRES-GFP plasmid^22^. All the PIEZO2 truncated versions were obtained from PIEZO2-GL by PCR Site directed mutagenesis, using KAPA HIFI PCR Kit (Roche; 07958838001) and primers (Integrated DNA Technologies) with common overhangs that mediate the recombination and and plasmid circularization inside the bacteria. The PIEZO1 truncated versions were also generated by PCR amplification of Piezo1-HA-IRES-GFP plasmid, which was generated by subcloning an HA sequence into a Piezo1-IRES-GFP plasmid^1^ using the previously described strategy. PIEZO2 and PIEZO1 PCR products, were digested with DpnI and directly transformed into XL10-Gold ultracompetent cells (200315; Agilent). For PIEZO2 mutants, bacteria were grown for 48 hours at 30°C while for PIEZO1 for 24 at 37°C respectively. For each mutant, several clones were screened by sequencing the mutated region and then the selected clone was fully sequenced to check the absence of unintended mutations.

### Cell culture and transfection

Neuro2a-P1KO cells (gift from Gary R. Lewin, MaxDelbrueck-Center for Molecular Medicine, Berlin, Germany) were grown at 37°C and 5% CO2 in a 1:1 mixture of Opti-MEM + Glutamax (Thermo Fisher Scientific; 51985-026) and DMEM (Dulbecco’s Modified Eagle Medium) (Thermo Fisher Scientific; 21969-035) supplemented with 10% FBS (Thermo Fisher Scientific; 10270106), 0.5% L-Glutamine (Thermo Fisher Scientific; 11539876), and 1% Penicillin-Streptomycin (Thermo Fisher Scientific; 15140-122). Cells were seeded on 13 mm of diameter Poly-L-lysine (Sigma; 15140-122) coated cover slips (Patch-Clamp and Immunocytochemistry) or in 6-well plates (Western-Blot and Cycloheximide assay).

To transfect the Neuro2a-PIEZO1-KO cells, we used the PEI transfection reagent (Linear PEI25K, PolyScience; 23966-1). Briefly, 24 hours after seeding, a mixture of PEI and Plasmid DNA diluted in DPBS (Dulbecco Phosphate Buffered Saline) (Sigma; D8537) was added to the cell culture media to a final concentration of: 112.5 ng/mL of PEI and 0.6 µg/mL of DNA. For PIEZO2 mutant plasmids cells were used 48 hours after transfection. For PIEZO1 cells were used 24 hours after transfection.

### Protein expression assessment by western blot

#### Total extract obtention

Western blot cultures were seeded at 290,000 cells per well and transfected following the established protocol. After 48 hours, cells were harvested by gentle detachment from the Petri dish using repeated pipetting with PBS. The number of collected cells was estimated by measuring their optical density at 600 nm. This value was used to calculate the amount of lysis buffer applied to each sample, thereby normalizing the amount of protein in each extract. Cells were lysed in the corresponding amount of RIPA buffer (HEPES 50 mM, NaCl 140 mM, Glycerol 10%, 1% Triton X-100, 1 mM EDTA, 2 mM EGTA, 0.5% Deoxycholate (Sigma; D6750), 1:100 Protease Inhibitor Complete (Roche; 1183617000)), with the maximum volume being 200 µl. After cell resuspension, samples were incubated for 1 hour at room temperature under constant rotation on a turning wheel. Following complete lysis, protein extracts were centrifuged at 12,000 RPM for 30 minutes to remove cell debris. The resulting total extracts were transferred to clean Eppendorf tubes and stored at −20°C until SDS-polyacrylamide gel electrophoresis (SDS-PAGE).

#### SDS-PAGE and Western blotting

9 µl of each sample were mixed with the same volume of Sample Buffer [Trizma Base 1M, SDS 10%, Bromophenol-Blue 0.05%, Glycerol 50%, reactants purchased from Sigma] and heated for 5 min at 95 °C. Next, the samples were loaded in a 10% polyacrylamide gel (0,75 mm thickness) and the SDS-PAGE gel was run at 120V for 110 min. The gel was then blotted onto a 0,45 µm Nitrocellulose membrane (Amersham; 10600007) using a transfer buffer [30 mM Trizma Base, 190 mM Glycine, 20% Methanol], and a fixed voltage of 60V for 16 hours. To prevent unspecific antibody binding, we incubated the membranes in 2% skimmed milk-TBS-T for 30 min. The membranes were subsequently incubated ON at 4°C with primary antibodies: rabbit anti-GFP at 1/500 to detect PIEZO2 (Abclonal; AE011), rabbit anti-HA to detect PIEZO1 at 1/1000 (Sigma; H6908), or rabbit anti-ß-Tubulin as loading control at 1/10,000 (BioLegend; 802001). After washing the membranes with TBS-T for 15 min in constant agitation, we incubated the membranes with the secondary antibody anti-Rabbit HRP conjugated (Millipore; AP182P) at 1/10,000 (GFP and HA) or 1/80,000 (ß-Tubulin). All antibodies were diluted in 5% skimmed milk-TBS-T. Lastly, the membranes were revealed using ECL select (Amersham; RPN2235), following the manufacturer’s instructions. Quantification images were acquired with the iBright CL1500 (Thermo Fisher Scientific-Fisher; A44114).

### Protein stability assays

To assess the protein stability, we stopped the protein synthesis by adding the transcription elongation blocker Cycloheximide. Similarly to the general western blot assay, we seeded 290,000 cells per well and transfected them following the protocol detailed above. 48 hours post-transfection, we added Cycloheximide solution prepared in DMSO at a final concentration of 50 µg/ml (Sigma; 01810). We then collected cells as described in the western blot section at the following time points: just after adding the Cycloheximide (T_0_) and after 4, 8, 12 and 24 hours. During these periods cells did not show major morphological alterations indicating that the times assayed cells were healthy. The collected samples were then processed as described in the western blot section previously described.

### Patch-clamp recordings of mechanically activated currents

Poking activated currents were recorded at room temperature in the patch-clamp whole cell mode. Patch pipettes with a tip resistance of 4–7MΩ were pulled with a flaming-Brown puller P-97 (Sutter Instruments) from borosilicate glass capillaries (Sutter Instruments; BF150-86-10) and filled with intracellular solution [125 mM K-gluconate, 7 mM KCl, 1 mM CaCl2, 1 mM MgCl2, 10 mM HEPES, 4 mM EGTA adjusted to pH 7.3 with KOH and 300 mOsm]. Cells were placed in a bath containing Extracellular solution [140 mM NaCl, 4 mM KCl, 2 mM CaCl2, 1 mM MgCl2, 4 mM glucose, and 10 mM HEPES, adjusted to pH 7.4 with NaOH and 290 mOsm]. Recordings were made with an EPC-10-USB double patch-clamp amplifier (HEKA) in combination with PATCHMASTER-NEXT (v1.3, HEKA). Pipette and membrane capacitances were compensated using the automatic function of PATCHMASTER-NEXT. Cells were clamped to a holding potential of −60 mV and mechanically stimulated by applying a train of stimuli increasing by 0.5 µm each at a standard stimulation velocity of 1 µm/ms (Changed only during the PIEZO2 velocity sensitivity experiment) by a piezo-driven micromanipulator (Nanomotor MM3A; Kleindiek Nanotechnik). The mechanical stimuli described were applied with a fired polished glass pipette (tip diameter 2-3 µm) positioned at a 45° angle relative to the cell surface, and currents were recorded with a sampling frequency of 50 kHz and filtered with a 2.9 kHz low-pass filter.

The inactivation time constant (Tinact) of the mechanically evoked currents was determined by fitting them to a single exponential function.

The mechanical thresholds of the mechanically evoked currents were determined by measuring the latency between the onset of the mechanical stimulus and the onset of the mechanically activated current. Mechanical activated currents were considered for analysis when the recorded current intensity was 10 times above the Standard Deviation from the baseline current. To determine the mechanical stimulus threshold, we proceeded as described in Verkest et al^23^. Briefly, we extracted the time form the stimuli trigger till the current overpass the 10*SD threshold. Then calculated the probe displacement considering the velocity of the probe. To test the PIEZO2 velocity sensitivity, we stimulate each cell with the standard indentation velocity (1 µm/ms) up to 5 µm, and then we apply the two different stimulation velocities (0.5 and 1.5 µm/ms). All currents were recorded with the same acquisition parameters and patch solutions described above.

Stretch activated and single channel currents were recorded at room temperature in the cell attached configuration. Currents were acquired with a sampling frequency of 50 kHz and filtered with a 2.9 kHz low-pass filter. Patch pipettes with a tip resistance of 5–8MΩ, covered with Sylgard (101697;Farnell) were pulled (Flaming-Brown puller P-97; Sutter Instruments) from borosilicate glass capillaries (Sutter Instruments, BF150-86-10), filled with intracellular solution [130 mM NaCl, 5 mM KCl, 1 mM CaCl2, 1 mM MgCl2, 10 mM TEA-Cl, and 10 mM HEPES, adjusted to pH 7.3 with NaOH and 300 mOsm]. Cells were placed in an extracellular solution [140 mM KCl, 1 mM CaCl2, 1 mM MgCl2, 10 mM glucose, and 10 mM HEPES, adjusted to pH 7.3 with KOH and 290 mOsm]. To elicit stretch activated currents, a 3 second pressure stimuli of different intensity were applied through the recording pipet with a High-Speed Pressure Clamp-1 (HPSC-1; ALA Scientific Instruments). To determine the unitary current of the channels, we applied a negative pressure stimulus of −60 mmHg during an I-V curve going from −100 mV to 0 mV in steps of 20 mV. Using a custom Python script, we calculated the currents as the difference between the background and the open state peaks of the Gaussian fit from the trace histogram. The unitary conductance then was calculated from the linear regression fit of the I-V of the individual cells. To determine the dose-response curve, we clamped the cells at −60 mV and applied a train of negative pressure pulses ranging from 0 to −60 mmHg in steps of −5 mmHg. We then estimate the charge transfer (Area Under the Curve, AUC), with a custom Python script over the whole stimulus in Pico Coulomb (pC) to quantify the response variations. These values were normalized to the maximal charge transfer and fitted to a Boltzmann distribution.

### Immunocytochemistry

For studying PIEZO subcellular distribution, N2a-P1-KO cells were co-transfected with one of the PIEZO constructs (PIEZO2-GL (wildtype and truncated version), PIEZO1-HA (Wild-Type and truncated versions) or AQP3-tGFF and AQP3-tGFP-P2-THU3) and one of the following intracellular compartment markers: ER-mCherry (Endoplasmic Reticulum) or Giantin-mCherry (cis-medial Golgi). For chemical membrane stain, cells were live-treated with Wheat Germ Agglutinin–Alexa Fluor 647 conjugate (Thermo Fisher Scientific; W32466) for 5 min on ice to reduce the internalization. Cells were washed 2 times for 10 min before continuing the standard immunocytochemistry protocol. Briefly, cells were fixed with 4% Paraformaldehyde (PFA) for 10 min, washed 3 more times, and permeabilized in a solution containing [2% donkey serum, 1% BSA, 0.1% Triton X-100, and 0.05% Tween-20] in PBS for 1 hour at RT. Samples were then incubated ON at 4°C with a dilution of 1/2,000 anti-GFP chicken antibody (Aves Lab; GFP-1020), or a dilution of 1/1,000 anti-turboGFP chicken antibody (Origene; TA150075) respectively, and a dilution of 1/1,000 anti-RFP rabbit antibody (Rockland; 600-401-379). After three washes with PBS for 10 min, cells were incubated for 1 hour at RT with a 1/1000 dilution of secondary antibodies containing AlexaFluor-488 donkey anti-chicken (Jackson InmunoResearch; 703-545-155) and AlexaFluor-568 donkey anti-rabbit (Invitrogen; A10042). All antibody dilutions were made in 1% BSA-PBS, and after three washes with PBS for 10 min, cells were placed in glass slides using Fluoro-Gel II with DAPI (Electron Microscopy Sciences; 17985-51). Cells transfected with P1-HA-IRES-GFP (Wild-Type or any of the truncated version) followed the same staining and mounting protocol described above but were incubated ON with a dilution of 1/2,000 anti-HA rabbit antibody (Sigma; H6908), and a 1/1000 dilution of AlexaFluor-488 donkey anti-chicken (Jackson InmunoResearch; 703-545-155).

### Image acquisition and analysis

Super-resolution images were acquired using an LSM880-AiryScan microscope (Zeiss) at 0.16 µm sections. Using a 63X/1.4 oil objective with a resolution limit of 87 nm. Acquired images were then deconvoluted using Huygens Professional Software (v23.1, Scientific Volume Imaging). The displayed images are z projections obtained from the acquired image set for each individual cell.

Person colocalization index and maps were also computed using Huygens Professional Software all images were thresholded using the gaussian minimum method.

Cluster size analysis was done with Imaris Software (v9.3.1, BitPlane). All surfaces were rendered using the Imaris surface tool setting the following parameters and filters: Surface detail of 0.1 µm; background subtraction option as thresholding method, this method applies a Gaussian filter (width 0.15 µm) to estimate the background value of each voxel, this value was then subtracted from every voxel in the image; finally, we applied a second filter (seed point diameter 0.15 µm) to split potential touching objects and ensure that we are analyzing individual objects.

### Data analysis and statistics

Electrophysiological recordings of mechanical evoked currents were analyzed using custom Python scripts. Super-Resolution images were analyzed using Huygens Professional Software (v23.1, Scientific Volume Imaging) and Imaris Software (v9.3.1, BitPlane). Western Blot images were analyzed using Fiji image analysis package.

Graphical data representation and statistical analysis was performed with Prism (v10, GraphPad). Data normality was first assessed using the Kolmogorov-Smirnov test and parametric or non-parametric tests were chosen accordingly. Statistical tests used thorough the study consist on t-test, one-way and two-way ANOVA, exact P-values, and the number of independent biological replicates are reported throughout the article. P-values are presented with results: both as numbers in the figure legends and as symbols (*P<0.05, **P<0.01, ***P<0.001, ns (not significant) P>0.05) in the graphs.

## Acknowledgements

This study was supported by the grant PID2020-116381GA-I00 funded by MCIN/AEI/ 10.13039/501100011033 and the grant CIDEGENT/2020/052 funded by the Generalitat Valenciana.

## Author contributions

S.S-F. cloned mutants, performed and analyzed patch-clamp recordings, immunocytochemistry and protein expression experiments and wrote the paper. E.S. cloned PIEZO2 mutants. F.J.T. conceptualized the study, acquired funding, analyzed the data, supervised the project and wrote the manuscript.

## Notes

CONFLICT OF INTEREST STATEMENT: The authors have declared that no conflict of interest exists.

### Competing Interest Statement

The authors have declared no competing interest.

## References

1. Coste, B. et al. Piezo1 and Piezo2 Are Essential Components of Distinct Mechanically Activated Cation Channels. Science 330, 55–60 (2010).

2. Murthy, S. E., Dubin, A. E. & Patapoutian, A. Piezos thrive under pressure: mechanically activated ion channels in health and disease. Nat. Rev. Mol. Cell Biol. 18, 771–783 (2017).

3. Wu, J., Lewis, A. H. & Grandl, J. Touch, Tension, and Transduction – The Function and Regulation of Piezo Ion Channels. Trends Biochem. Sci. 42, 57–71 (2017).

4. Holt, J. R. et al. Spatiotemporal dynamics of PIEZO1 localization controls keratinocyte migration during wound healing. eLife 10, e65415 (2021).

5. Schrenk-Siemens, K. et al. PIEZO2 is required for mechanotransduction in human stem cell–derived touch receptors. Nat. Neurosci. 18, 10–16 (2015).

6. Florez-Paz, D., Bali, K. K., Kuner, R. & Gomis, A. A critical role for Piezo2 channels in the mechanotransduction of mouse proprioceptive neurons. Sci. Rep. 6, 25923 (2016).

7. Woo, S.-H. et al. Piezo2 is the principal mechanotransduction channel for proprioception. Nat. Neurosci. 18, 1756–1762 (2015).

8. Woo, S.-H. et al. Piezo2 is required for Merkel-cell mechanotransduction. Nature 509, 622–626 (2014).

9. Chesler, A. T. et al. The Role of *PIEZO2* in Human Mechanosensation. N. Engl. J. Med. 375, 1355–1364 (2016).

10. Ranade, S. S. et al. Piezo2 is the major transducer of mechanical forces for touch sensation in mice. Nature 516, 121–125 (2014).

11. Nonomura, K. et al. Mechanically activated ion channel PIEZO1 is required for lymphatic valve formation. Proc. Natl. Acad. Sci. 115, 12817–12822 (2018).

12. Lam, R. M. et al. PIEZO2 and perineal mechanosensation are essential for sexual function. Science 381, 906–910 (2023).

13. Marshall, K. L. et al. PIEZO2 in sensory neurons and urothelial cells coordinates urination. Nature 588, 290–295 (2020).

14. Prato, V. et al. Functional and Molecular Characterization of Mechanoinsensitive “Silent” Nociceptors. Cell Rep. 21, 3102–3115 (2017).

15. Fernández-Trillo, J. et al. Piezo2 Mediates Low-Threshold Mechanically Evoked Pain in the Cornea. J. Neurosci. 40, 8976–8993 (2020).

16. Szczot, M. et al. PIEZO2 mediates injury-induced tactile pain in mice and humans. Sci. Transl. Med. 10, eaat9892 (2018).

17. Yang, X. et al. Structure deformation and curvature sensing of PIEZO1 in lipid membranes. Nature 604, 377–383 (2022).

18. Guo, Y. R. & MacKinnon, R. Structure-based membrane dome mechanism for Piezo mechanosensitivity. eLife 6, (2017).

19. Zhao, Q. et al. Structure and mechanogating mechanism of the Piezo1 channel. Nature 554, 487–492 (2018).

20. Saotome, K. et al. Structure of the mechanically activated ion channel Piezo1. Nature 554, 481–486 (2018).

21. Wang, L. et al. Structure and mechanogating of the mammalian tactile channel PIEZO2. Nature 573, 225–229 (2019).

22. Taberner, F. J. et al. Structure-guided examination of the mechanogating mechanism of PIEZO2. Proc. Natl. Acad. Sci. 201905985 (2019) doi:10.1073/pnas.1905985116.

23. Verkest, C. et al. Intrinsically disordered intracellular domains control key features of the mechanically-gated ion channel PIEZO2. Nat. Commun. 13, 1365 (2022).

24. Moroni, M., Servin-Vences, M. R., Fleischer, R., Sánchez-Carranza, O. & Lewin, G. R. Voltage gating of mechanosensitive PIEZO channels. Nat. Commun. 9, 1096 (2018).

25. Mulhall, E. M. et al. Direct observation of the conformational states of PIEZO1. Nature 620, 1117–1125 (2023).

26. Ridone, P. et al. Disruption of membrane cholesterol organization impairs the activity of PIEZO1 channel clusters. J. Gen. Physiol. 152, e201912515 (2020).

27. Shaw, J. L., Pablo, J. L. & Greka, A. Mechanisms of Protein Trafficking and Quality Control in the Kidney and Beyond. Annu. Rev. Physiol. 85, 407–423 (2023).

28. Soler, D. C., Ballesteros, A., Sloan, A. E., McCormick, T. S. & Stepanyan, R. Multiple plasma membrane reporters discern LHFPL5 region that blocks trafficking to the plasma membrane. Sci. Rep. 13, 2528 (2023).

29. Bannykh, S. I., Rowe, T. & Balch, W. E. The organization of endoplasmic reticulum export complexes. J. Cell Biol. 135, 19–35 (1996).

30. Hughes, H. et al. Organisation of human ER-exit sites: requirements for the localisation of Sec16 to transitional ER. J. Cell Sci. 122, 2924–2934 (2009).

31. Sanz-Salvador, L., Andrés-Borderia, A., Ferrer-Montiel, A. & Planells-Cases, R. Agonist- and Ca2+-dependent Desensitization of TRPV1 Channel Targets the Receptor to Lysosomes for Degradation. J. Biol. Chem. 287, 19462–19471 (2012).

32. Paillart, C. et al. ACTIVITY-INDUCED INTERNALIZATION AND RAPID DEGRADATION OF SODIUM CHANNELS IN CULTURED FETAL NEURONS. Biol. Cell 86, 187–187 (1996).

33. Gong, Q., Keeney, D. R., Molinari, M. & Zhou, Z. Degradation of Trafficking-defective Long QT Syndrome Type II Mutant Channels by the Ubiquitin-Proteasome Pathway. J. Biol. Chem. 280, 19419–19425 (2005).

34. Zhou, Z., Gong, Q. & January, C. T. Correction of Defective Protein Trafficking of a Mutant HERG Potassium Channel in Human Long QT Syndrome. J. Biol. Chem. 274, 31123–31126 (1999).

35. Krshnan, L., Van De Weijer, M. L. & Carvalho, P. Endoplasmic Reticulum– Associated Protein Degradation. Cold Spring Harb. Perspect. Biol. a041247 (2022) doi:10.1101/cshperspect.a041247.

36. Ciardo, M. G. et al. Whirlin increases TRPV1 channel expression and cellular stability. Biochim. Biophys. Acta BBA - Mol. Cell Res. 1863, 115–127 (2016).

37. Glogowska, E. et al. Novel mechanisms of PIEZO1 dysfunction in hereditary xerocytosis. Blood 130, 1845–1856 (2017).

38. Ma, S. et al. Common PIEZO1 Allele in African Populations Causes RBC Dehydration and Attenuates Plasmodium Infection. Cell 173, 443–455.e12 (2018).

39. Coste, B. et al. Gain-of-function mutations in the mechanically activated ion channel PIEZO2 cause a subtype of Distal Arthrogryposis. Proc. Natl. Acad. Sci. 110, 4667–4672 (2013).

40. Ma, S. et al. Excessive mechanotransduction in sensory neurons causes joint contractures. Science 379, 201–206 (2023).

41. Wu, J., Goyal, R. & Grandl, J. Localized force application reveals mechanically sensitive domains of Piezo1. Nat. Commun. 7, 12939 (2016).

42. Soler, D. C. et al. An uncharacterized region within the N-terminus of mouse TMC1 precludes trafficking to plasma membrane in a heterologous cell line. Sci. Rep. 9, 15263 (2019).

43. Campbell, B. C. et al. mGreenLantern: a bright monomeric fluorescent protein with rapid expression and cell filling properties for neuronal imaging. Proc. Natl. Acad. Sci. U. S. A. 117, 30710–30721 (2020).

